# Conserved yet Divergent Smc5/6 Complex Degradation by Mammalian Hepatitis B Virus X Proteins

**DOI:** 10.1101/2024.10.28.620692

**Authors:** Maya Shofa, Yuri V Fukushima, Akatsuki Saito

**Affiliations:** Department of Veterinary Science, Faculty of Agriculture, University of Miyazaki, Miyazaki, Miyazaki 8892192, Japan; Graduate School of Medicine and Veterinary Medicine, University of Miyazaki, Miyazaki, Miyazaki 8891692, Japan; Center for Animal Disease Control, University of Miyazaki, Miyazaki, Miyazaki 8892192, Japan

**Keywords:** hepatitis B virus (HBV), domestic cat HBV (DCHBV), structural maintenance of chromosome (Smc) Smc5/6 complex, DNA-binding protein 1 (DDB1)

## Abstract

Hepatitis B virus (HBV), belonging to the genus *Orthohepadnavirus*, can cause chronic hepatitis and hepatocarcinoma in humans. HBV ensures optimal replication by encoding X, a multifunctional protein responsible for degrading the structural maintenance of the chromosome (Smc) 5/6 complex, an anti-HBV factor in hepatocytes. Previous studies suggest that the degradation activity of the Smc5/6 complex is conserved among viruses from the genus *Orthohepadnavirus*. Recently, a novel hepadnavirus in cats, domestic cat hepadnavirus (DCH) or DCH B virus (DCHBV), has been identified to be genetically close to HBV. However, it remains unclear whether the DCHBV X protein possesses a similar Smc5/6 complex–degrading activity.

Here, we investigated the degradation activity of the Smc5/6 complex by X of the viruses of the genus *Orthohepadnavirus*, including DCHBV, in cells derived from primates and cats. We found that the DCHBV X protein degraded Smc6 in the cells of several host species, and the degree of its anti-Smc5/6 activity differed depending on the host species. Furthermore, we demonstrated that the DCHBV X protein degraded Smc6 independently of DNA-binding protein 1 (DDB1), which is a critical host factor for HBV X–mediated Smc6 degradation. Our findings highlight the conserved yet divergent degradation machinery of the Smc5/6 complex of mammalian hepatitis B virus X proteins.

**Importance:** Hepatitis B virus (HBV) causes chronic hepatitis and liver cancer in humans. HBV mainly acts by degrading the Smc5/6 complex, a defense mechanism in liver cells that restricts viral replication. Smc5/6 degradation, mediated by the HBV protein X, is conserved across the viruses in the same genus. However, it remains unclear whether the novel domestic cat HBV (DCHBV) exhibits similar capabilities.

Here, we compared Smc5/6 degradation by the X proteins of various HBVs, including DCHBV, in different primate and feline cells. The DCHBV X protein could degrade Smc6 in various host cells at different levels. Notably, DCHBV X could degrade Smc6 without requiring DDB1, a host factor essential for HBV X–mediated Smc6 degradation. These findings highlight the conserved yet distinct strategies employed by mammalian HBVs to evade the host Smc5/6 system, expanding our understanding of viral adaptation and persistence across species.

## Introduction

Hepatitis B virus (HBV), of the family *Hepadnaviridae* family and genus *Orthohepadnavirus*, is an enveloped virus with a partially double-stranded, circular DNA genome. Approximately 250 million individuals are chronically infected with HBV worldwide, and chronic HBV infections can cause cirrhosis and hepatocellular carcinoma (1). Several therapeutic agents, including nucleoside analogs and interferon-α (IFN-α), have been approved to treat HBV infection; however, curing the infection, i.e., eliminating the virus from an individual, remains challenging. Therefore, there has been much research on developing antiviral therapies to achieve a cure. In addition, there is a need to identify new animal models for studying HBV infection and to study the interaction between host defense mechanisms and HBV. Therefore, it is important to elucidate the molecular interactions between hosts and viruses.

The viruses belonging to the genus *Orthohepadnavirus* encode 4 viral proteins: polymerase (P) protein, surface (S) protein, core (C) protein, and X protein (17). While S and C proteins are structural proteins, the P protein is responsible for viral genome replication (18). Meanwhile, the HBV X protein is a versatile regulator that influences host cellular processes and activates multiple transcriptional factors, including AP-1, AP-2, NF-κB, and the cAMP response element (reviewed in 19), playing a critical role in HBV replication and pathogenesis (20). The HBV X protein comprises 4 key regions that are essential for transactivation, dimerization, p53 interaction, and binding to the 14-3-3 protein motif (reviewed in 21). Notably, HBV X protein antagonizes the inhibitory effect by the structural maintenance of the chromosome (Smc) Smc5/6 complex on HBV in infected cells. HBV X protein acts by degrading the Smc5/6 complex via interaction with the DNA-binding protein 1 (DDB1)–Cul4 ubiquitin ligase machinery (22–24). Moreover, the genus *Orthohepadnavirus* also includes HBV-like viruses that infect mammals, such as rodents, bats, and primates, and the function of X from viruses in the genus *Orthohepadnavirus* is conserved in primates, bats, and rodents (25).

However, it is unclear if the function of X is conserved in HBVs found in other animals. For example, domestic cat hepadnavirus (DCH), an HBV-like virus also classified as domestic cat hepatitis B virus (DCHBV), was identified in domestic cats in 2018. DCHBV infection displayed a signature of chronic hepatitis, suggesting an association between them (2). DCHBV has been identified in several countries and regions, including Australia (3, 4), Italy (5), the UK (6), the US, Malaysia (7), Thailand (8), Taiwan (9), Türkiye (10), Japan (11, 12), Hong Kong (13), Chile (14), and Brazil (15). Furthermore, we have recently demonstrated that DCHBV shares with HBV the sodium/bile acid cotransporter (NTCP) as an entry receptor (16); this finding suggests that DCHBV infection in cats can be a promising animal model for studying HBV infection. However, whether the function of the DCHBV X protein is conserved in carnivores remains to be elucidated.

In this study, we analyzed the degradation of the Smc5/6 complex by mammalian hepatitis B virus X proteins in primate and feline cells. While Western blotting is used to study Smc5/6 degradation by mammalian hepatitis B virus X proteins, this assay has a relatively low throughput. Thus, we improved the assay’s throughput by developing a new assay using a split-type red fluorescent protein to measure Smc5/6 degradation by mammalian hepatitis B virus X proteins quantitively. We observed that while the human–derived HBV(A) X protein showed conserved Smc6 degradation activity in primate and feline cells, the DCHBV X protein displayed weaker Smc6 degradation activity in the cells of African green monkeys. Furthermore, we uncovered that the DCHBV(KT116) X protein degraded the Smc5/6 complex via a different mechanism compared with the HBV(A) X protein.

Our findings suggest that the X proteins in the viruses in the genus *Orthohepadnavirus* may have evolved to have conserved but differential Smc5/6 degradation activities, depending on the combination of viruses and host species. Collectively, our findings highlight the evolutionary traits of the mammalian hepatitis B virus X proteins as a consequence of the arms race between hosts and viruses.

## Results

### Genetic similarity of the mammalian hepatitis B virus X proteins

First, we analyzed the genetic diversity of mammalian hepatitis B virus X proteins by aligning the sequences of the X proteins of 5 strains of HBV, 5 strains of DCHBV, 5 species of bat hepadnavirus, 1 strain of woodchuck hepatitis virus, and those of other viruses belonging to the genus *Orthohepadnavirus* (**Figure 1A**). Interestingly, DCHBV clustered independently but closely with domestic donkey HBV and bat HBV. Additionally, DCHBV exhibited distinct clustering patterns compared with primate HBV **(Figure 1B).**

**Figure 1.**
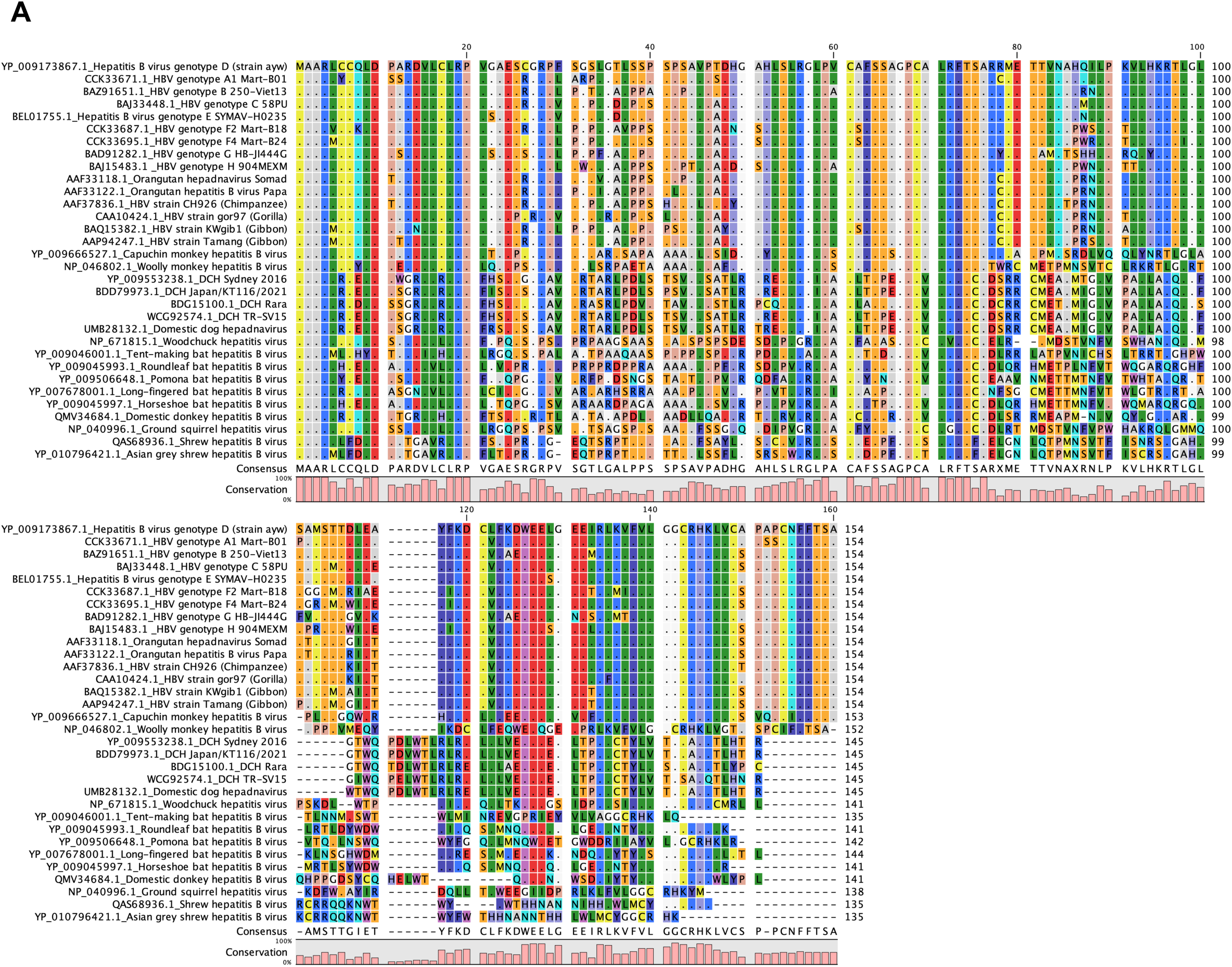
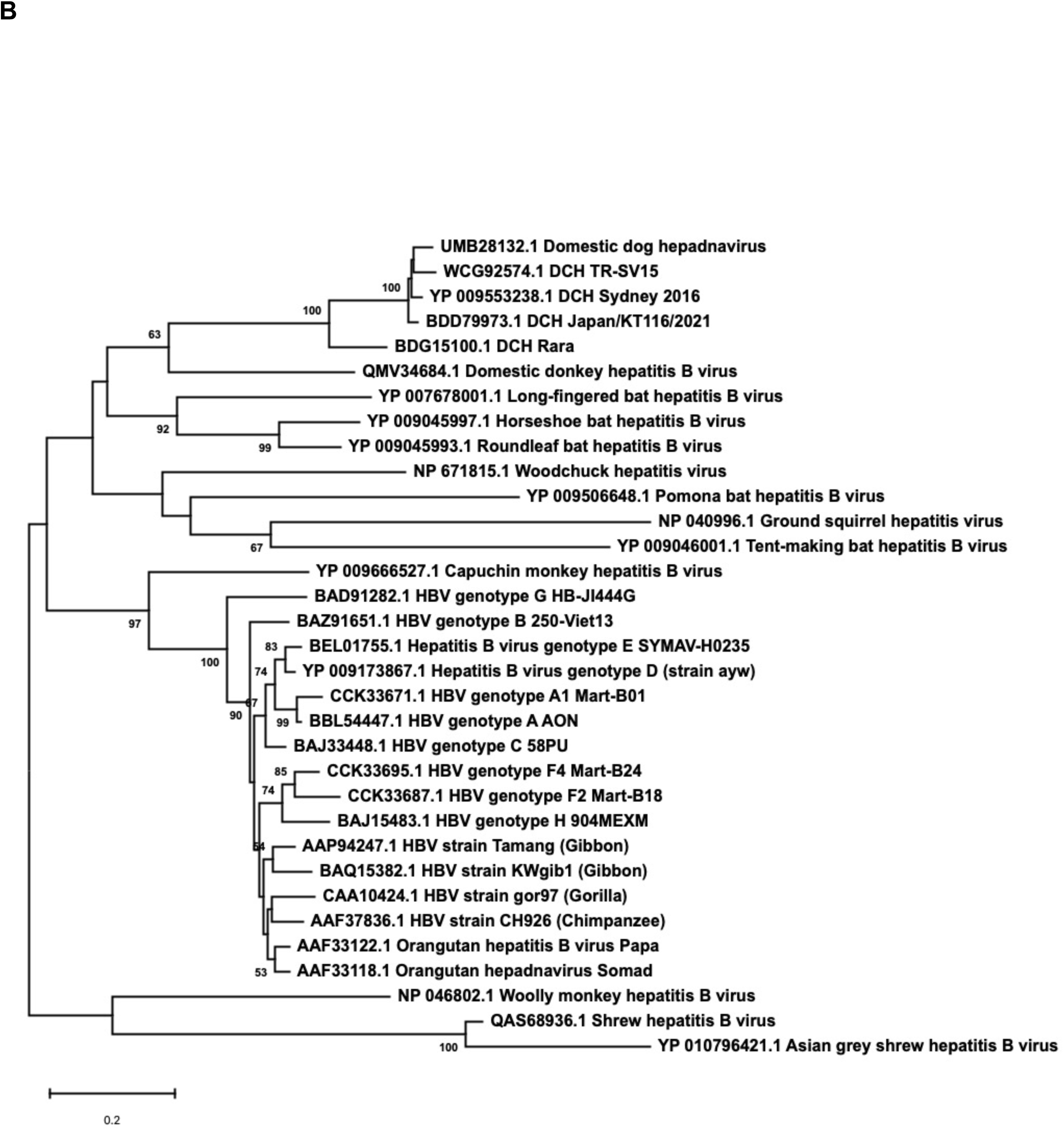
Sequence alignment and phylogenetic analysis of the X proteins from viruses of the genus *Orthohepadnavirus*. (A) Amino acid sequence alignment of the X proteins of the genus *Orthohepadnavirus*. The residue numbering was based on the HBV genotype D strain aw. (B) The phylogenetic tree was constructed using the alignment of X protein sequences derived from viruses of the genus *Orthohepadnavirus*. The evolutionary history was inferred using the neighbor-joining method. The percentage of replicate trees in which the associated taxa clustered together in the bootstrap test (1000 replicates) are shown next to the branches.

### Similarity in Smc6 among mammals

Next, we analyzed the similarities in Smc6 between humans and other mammals, including cats (**Figure 2**). According to the phylogenetic tree, feline Smc6 was genetically close to primate Smc6 even though they were on different branches. The similarity rates of mammalian Smc6 to human Smc6 at the amino acid level were consistently high, with values of more than 90%, indicating relative conservation across species. Notably, platypus showed a lower sequence identity, at less than 70%, with human Smc6, suggesting greater evolutionary divergence in this species (**Supplementary Table 1**). Also, we found a 5-amino-acid indel in the N-terminal region of platypus Smc6 (**Figure 2**). Unlike hominoids and Old World monkeys (OWMs), New World monkeys and other mammals possessed a TXSFX motif between positions 36 and 48. Furthermore, feline Smc6 (isoform 1) had an additional 5-amino-acid insertion at positions 216-KVRNT-222, which is absent in feline Smc6 (isoform 2) and other mammals (**Figure 2**).

**Figure 2.**
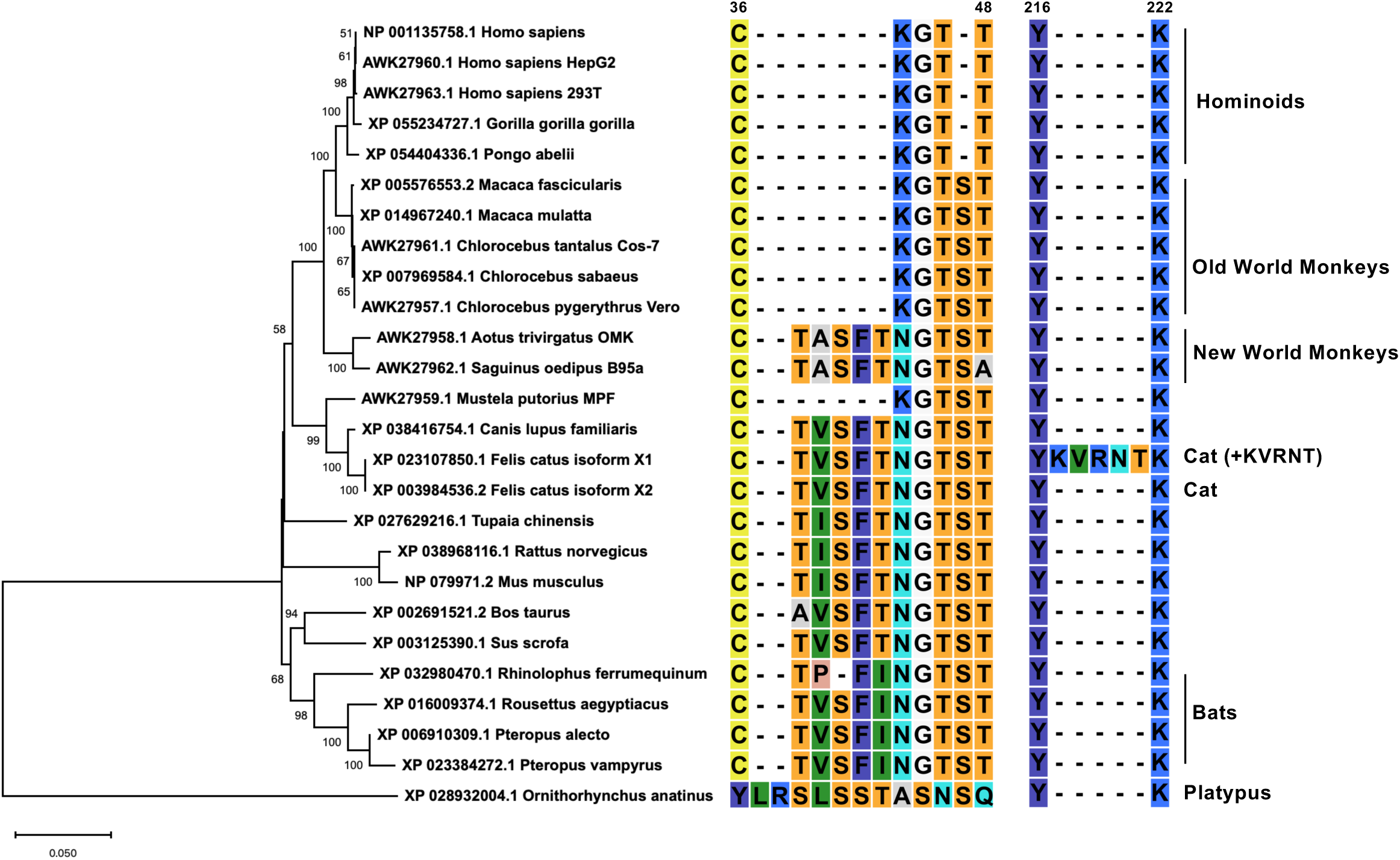
Sequence alignment of mammalian Smc6 and phylogenetic analysis of the entire mammalian Smc6. The alignment of the amino acid sequence of mammalian Smc6, specifically amino acids 36 to 48 and 216 to 222, was conducted. Subsequently, a phylogenetic tree for the complete mammalian Smc6 was constructed. The numbering of residues was standardized in reference to the human Smc6 sequence.

### Mammalian hepatitis B virus X proteins exhibit differential Smc5/6 degradation activity depending on the host species

Four cell lines derived from the 3 mammalian species, i.e., Lenti-X 293T (human), COS-7 (African green monkey: AGM), Fcwf-4 (cat), and CRFK (cat), were transfected with expression vectors encoding mNeonGreen or mammalian hepatitis B virus X proteins tagged with mNeonGreen. We observed comparable levels of mNeonGreen-tagged X in these cell lines **(Figure 3, right side of each panel)**. While the expression of mNeonGreen alone did not affect the level of the Smc5/6 complex in each cell line, all mammalian hepatitis B virus X proteins showed a significant degradation activity of the Smc5/6 complex in Lenti-X 293T and Fcwf-4 cells **(Figures 3A and 3D)**. In contrast, we found a differential Smc5/6 degradation activity of mammalian hepatitis B virus X proteins in COS-7 and CRFK cells **(Figure 3B and 3D)**. Specifically, HBV(A) X had a significant Smc5/6 degradation activity in COS-7 and CRFK cells. This result was consistent with a previous finding that HBV(A) X has a conserved degradation activity with all mammalian Smc6, including in cell lines derived from OWMs (Vero cells and COS-7 cells) (25). While DCHBV(KT116) X efficiently degraded the Smc5/6 complex in Lenti-X 293T, Fcwf-4, and CRFK cells **(Figures 3A, 3C, and 3D)**, it caused minimal Smc5/6 degradation in COS-7 cells **(Figure 3B)**. In CRFK cells, HBV(H) X and orangutan HBV (OHBV) X failed to degrade Smc6 **(Figure 3D)**. These results suggest that mammalian hepatitis B virus X proteins have differential Smc6 degradation activities depending on the host species and cell types.

**Figure 3.**
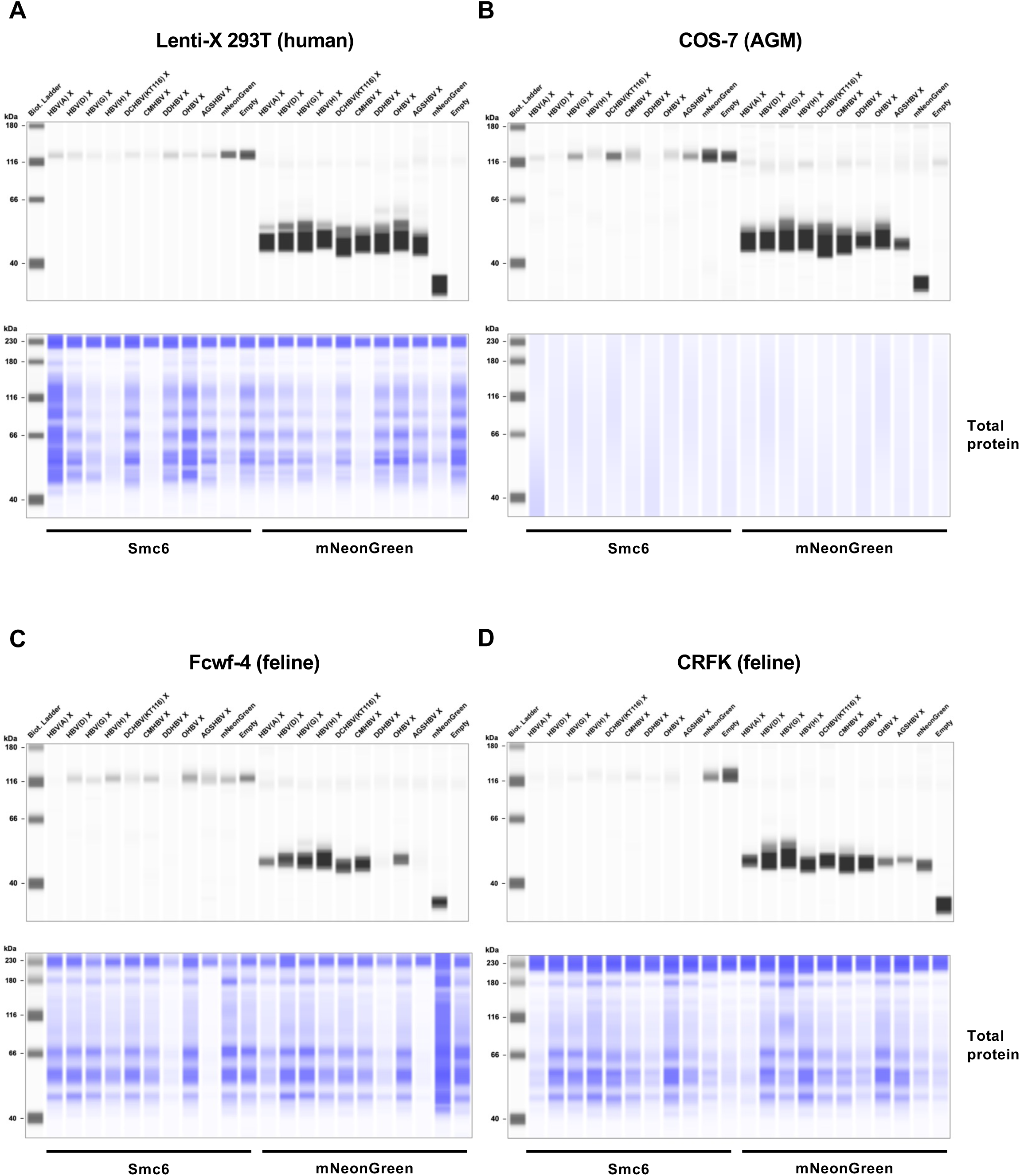
Mammalian hepatitis B virus X proteins exhibit differential Smc5/6 complex degradation activities depending on the host species. Lenti-X 293T (human), COS-7 (African green monkey), Fcwf-4 (feline), and CRFK (feline) cells were transfected with plasmids encoding mNeonGreen-HBV(A) X, mNeonGreen-DCHBV(KT116) X, or mNeonGreen or only pCAGGS. The levels of Smc6 and mNeonGreen at 2 days after transfection were analyzed by Western blotting. The results are representative of at least 3 independent experiments.

### DCHBV(KT116) X is more predominantly localized in the nucleus than HBV(A) X

While the subcellular localization of HBV X is both cytoplasmic and nuclear (26–28), about 80% of it is concentrated in the cytoplasm (26). We examined the similarity of X subcellular localizations by comparing the localization of HBV(A) X and DCHBV(KT116) X in Lenti-X 293-T cells. While HBV X was observed both in the cytoplasm and nucleus, DCHBV(KT116) X was mainly localized in the nucleus **(Figure 4A)**. In addition, we evaluated the distribution of X using a subcellular fractionation assay. First, we confirmed the efficient separation of the cytosolic and nuclear fractions **(Figure 4B)**. Second, we observed that while HBV(A) X was distributed in both the cytoplasm and the nucleus, DCHBV(KT116) X was predominantly present in the nucleus **(Figure 4C)**; these findings are consistent with observations made with fluorescence microscopy **(Figure 4A)**.

**Figure 4.**
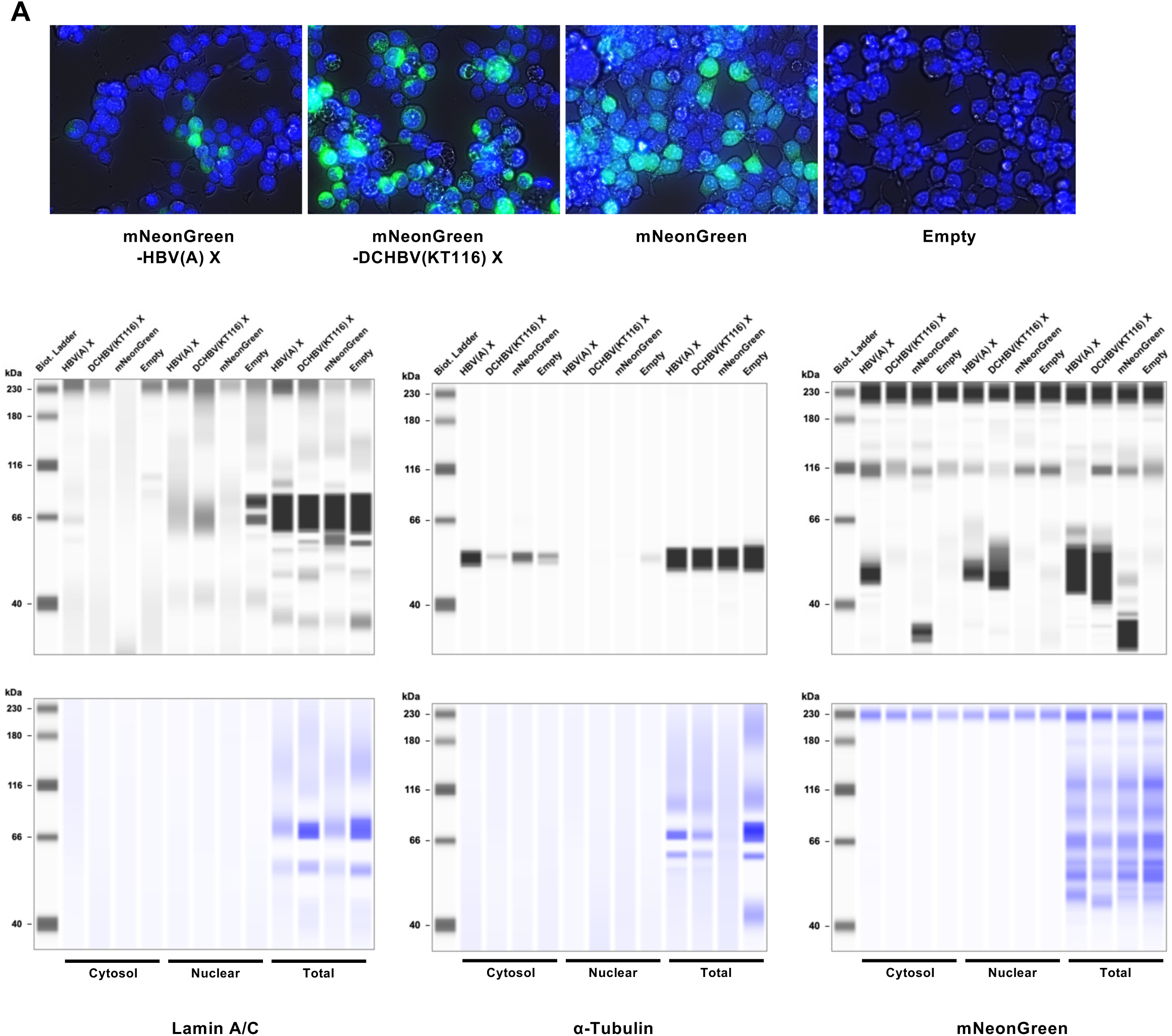
Subcellular localization of HBV(A) X and DCHBV(KT116) X. (A) Lenti-X 293-T cells were transfected with pCAGGS plasmids encoding mNeonGreen-HBV(A) X, mNeonGreen-DCHBV(KT116) X, mNeonGreen or only pCAGGS. (B) The nuclear fraction, cytosolic fraction, and whole-cell lysates (Total) were analyzed by Western blotting using anti-lamin A/C (nuclear), anti-α-tubulin (cytosolic), and anti-mNeonGreen (X protein) antibodies. The results are representative of at least 3 independent experiments.

### DCHBV(KT116) X degrades the Smc5/6 complex independently of DDB1

We improved the throughput of the analysis of Smc5/6 degradation activities by developing a new assay using a split-type of a red fluorescent protein, sfCherry3C (29). First, we tested the specificity of this system by transfecting Lenti-X 293T cells with either pCAGGS-SpyCatcher-sfCherry3C(1–10), pCAGGS-SpyTag-sfCherry3C(11)-mNeonGreen alone, or both. We found that pCAGGS-SpyCatcher-sfCherry3C(1–10) or pCAGGS-SpyTag-sfCherry3C(11)-mNeonGreen transfection did not produce any sfCherry3C signal. In contrast, the co-transfection of both plasmids produced double-positive signals of sfCherry3C and mNeonGreen (**Figure 5A**). We observed efficient levels of sfCherry3C after the co-transfection of pCAGGS-SpyCatcher-sfCherry3C(1–10) and pCAGGS-SpyTag-sfCherry3C(11)-Smc6 (Human) **(Figure 5A)**; the finding highlighted the specificity of this system **(Supplementary Figure 1)**.

**Figure 5.**
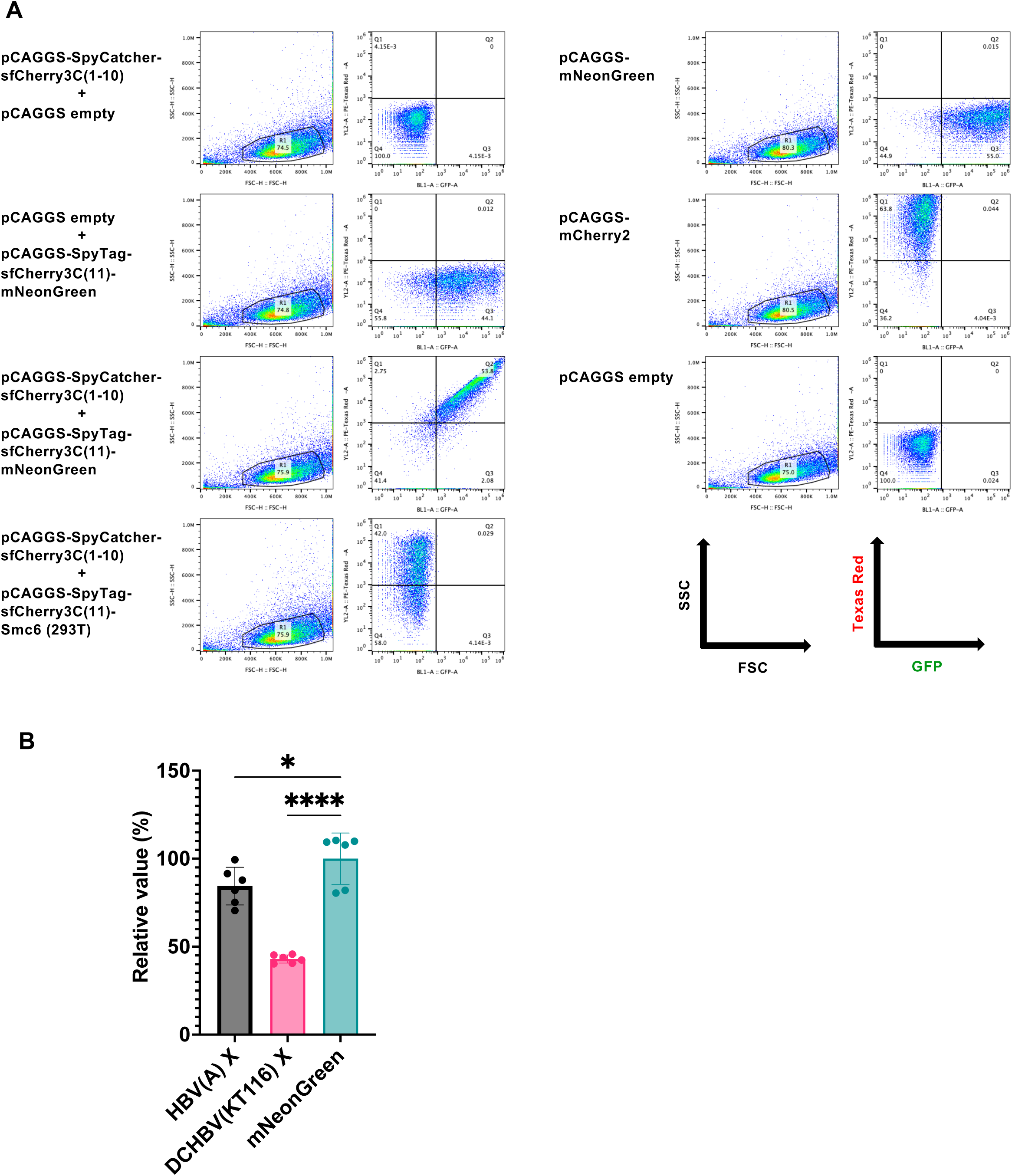
A flow cytometry–based SMC6 degradation assay using sfCherry3C-Smc6. (A) Lenti-X 293T cells were transfected with pCAGGS plasmids encoding SpyCatcher-sfCherry3C(1–10) only, SpyTag-sfCherry3C(11)-mNeonGreen only, or in combination of SpyCatcher-sfCherry3C(1–10), SpyTag-sfCherry3C(11)-mNeonGreen, and SpyTag-sfCherry3C(11)-HA-Smc6 (human). The signals of mNeonGreen and sfCherry3C were measured 2 days after transfection. (B) Lenti-X 293T cells were co-transfected with pCAGGS plasmids encoding SpyCatcher-sfCherry3C(1–10) only, SpyTag-sfCherry3C(11)-HA-Smc6 (human) only, or in combination with mNeonGreen-HBV(A) X, mNeonGreen-DCHBV(KT116) X, or mNeonGreen. Smc6 degradation was evaluated by measuring sfCherry3C levels using flow cytometry. The results, presented as the mean and SD of sextuplicate measurements from 1 assay, are representative of at least 3 independent experiments. The differences in sfCherry3C positivity between the cells producing HBV(A) X, DCHBV(KT116) X, and mNeonGreen were evaluated using one-way ANOVA, followed by the Tukey test. \**p* < 0.05, \*\*\**p* < 0.001, \*\*\*\**p* < 0.0001, ns (not significant).

Next, Lenti-X 293T cells were co-transfected with pCAGGS-SpyCatcher-sfCherry3C(1–10), pCAGGS-SpyTag-sfCherry3C(11)-Smc6 (Human, Cat, and Cat (+KVRNT)), and pCAGGS plasmids encoding mNeonGreen-tagged X proteins. The co-expression of the plasmids encoding HBV(A) X and DCHBV(KT116) X resulted in a lower level of sfCherry3C **(Figure 5C)**, suggesting that this system could reproduce the Smc6 degradation visualized by Western blotting **(Figure 3)**.

HBV X degrades the Smc5/6 complex by interacting with the host DDB1-containing E3 ubiquitin ligase complex for ubiquitin-mediated Smc5/6 degradation (22, 23). Thus, we determined whether HBV(A) X and DCHBV(KT116) X shared the Smc6 degradation mechanism by examining the effect of *Ddb1* deletion on Smc5/6 degradation (**Figure 6A**). While knockdown of human *Ddb1* in Lenti-X 293T cells completely inhibited HBV(A) X–mediated Smc5/6 degradation (**Figure 6B**), it did not affect DCHBV(KT116) X–induced Smc5/6 degradation (**Figure 6B**). This finding suggested that DCHBV(KT116) X degraded the Smc5/6 complex in a different mechanism from that of HBV(A) X.

**Figure 6.**
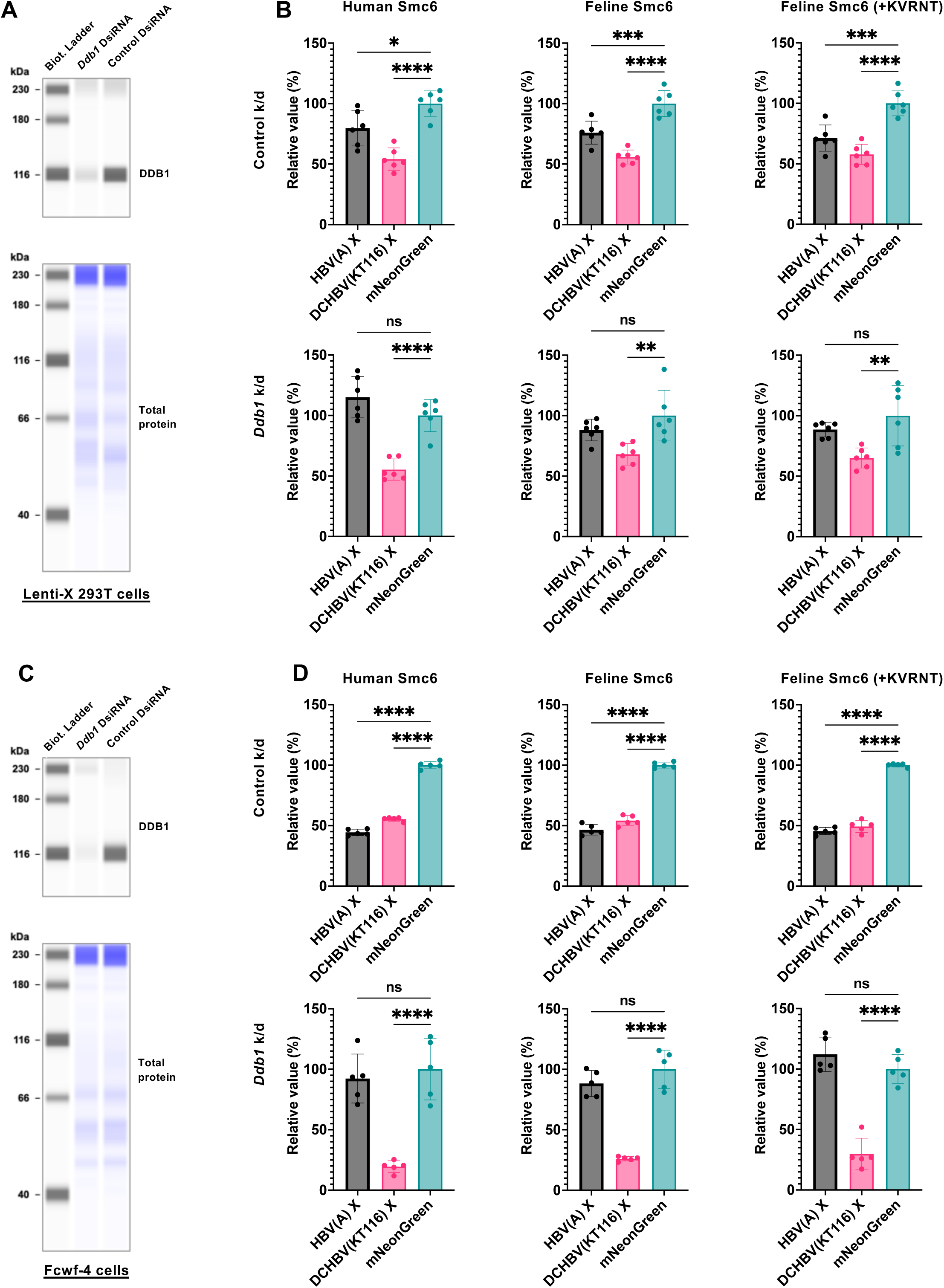
DCHBV(KT116) X degrades the Smc5/6 complex independently of DDB1. (A) Lenti-X 293-T cells were transfected with DsiRNA targeting human *Ddb1* or non-targeting control DsiRNA. The level of DDB1 in Lenti-X 293-T cells was determined using Western blotting. (B) After 2 days of DsiRNA transfection, the cells were co-transfected with pCAGGS plasmids encoding SpyCatcher-sfCherry3C(1–10), SpyTag-sfCherry3C(11)-HA-Smc6 (human, feline, or feline with KVRNT insertion), or in combination with mNeonGreen-HBV(A) X, mNeonGreen-DCHBV(KT116) X, or mNeonGreen. Smc6 degradation was evaluated by measuring sfCherry3C levels with flow cytometry. The results, presented as the mean and SD of sextuplicate measurements from 1 assay, are representative of at least 3 independent experiments. The differences in sfCherry3C positivity between the cells producing HBV(A) X, DCHBV(KT116) X, and mNeonGreen were evaluated using one-way ANOVA, followed by the Tukey test. \**p* < 0.05, \*\**p* < 0.01, \*\*\**p* < 0.001, \*\*\*\**p* < 0.0001, ns (not significant). (C) Fcwf-4 cells were transfected with DsiRNA targeting cat *Ddb1* or non-targeting control DsiRNA. The level of Ddb1 in Fcwf-4 cells was determined by Western blotting. (D) After 2 days of DsiRNA transfection, the cells were co-transfected with pCAGGS plasmids encoding SpyCatcher-sfCherry3C(1–10), SpyTag-sfCherry3C(11)-HA-Smc6 (human, feline, or feline with +KVRNT insertion), in combination with mNeonGreen-HBV(A) X, mNeonGreen-DCHBV(KT116) X, or mNeonGreen. Smc6 degradation was evaluated by measuring sfCherry3C levels with flow cytometry. The results, presented as the mean and SD of sextuplicate measurements from 1 assay, are representative of at least 3 independent experiments. The differences in sfCherry3C positivity between cells producing HBV(A) X, DCHBV(KT116) X, and mNeonGreen were evaluated by one-way ANOVA, followed by the Tukey test. \*\*\**p* < 0.001, \*\*\*\**p* < 0.0001, ns (not significant).

Further, we tested the role of feline DDB1 in X-mediated Smc6 degradation in feline-derived Fcwf-4 cells (**Figure 6C**). The knockout of *Ddb1* negated the Smc6 degradation induced by HBV(A) X (**Figures 6D**), suggesting a similar role for human and feline DDB1 in HBV(A) X-mediated Smc5/6 degradation. In contrast, Smc6 degradation by DCHBV(KT116) X was more efficient in *Ddb1*-depleted cells. These results suggest that human DDB1 and feline DDB1 play different roles in DCHBV(KT116) X–mediated Smc5/6 degradation.

## Discussion

In this study, we have demonstrated that the Smc5/6 degradation activities, mediated by mammalian hepatitis B virus X proteins, are divergent depending on the combination of the virus and the host species. Further, we have uncovered that HBV(A) X and DCHBV(KT116) X depend on DDB1 differently for Smc5/6 degradation.

The phylogenetic analysis of DCHBV X indicated its placement on a branch separate from that of HBV **(Figure 1)**. In addition, Smc6 was found to be divergent in mammals (**Figure 2**), suggesting different Smc5/6 degradation activities depending on the virus and host. Supporting this suggestion, the Western blotting analysis demonstrated that HBV(A) X degraded Smc6 in human, monkey, and feline cells. In contrast, DCHBV(KT116) X minimally degraded Smc6 in monkey cells **(Figures 3A, 3B, 3C, and 3D)**, suggesting that it has a narrower range of Smc5/6 degradation than HBV(A) X. In addition, several X proteins, including HBV(H) X and orangutan HBV X, minimally degraded Smc6 in feline cells **(Figure 3D)**, suggesting that the combination of the virus and host species significantly affects Smc5/6 degradation. Alternatively, feline cells may have the machinery to antagonize Smc6 degradation by several mammalian hepatitis B virus X proteins.

Using our novel Smc6 degradation assay (**Figure 5 and Supplementary Figure 1**), we demonstrated that HBV(A) X–mediated Smc6 degradation was negated by Ddb1 depletion (**Figure 6B**). This finding is consistent with previous reports that HBV(A) X requires interaction with DDB1 for Smc6 degradation (22, 23). In marked contrast, DCHBV(KT116) X degraded Smc6 independently of DDB1 (**Figure 6D**).

Another difference between HBV(A) X and DCHBV(KT116) X is their cellular localization. DCHBV(KT116) X protein was predominantly localized in the nucleus compared with HBV(A) X (**Figures 4A and 4C**). The cellular localization of HBV X affects the functions of X (28, 30–32). Therefore, the predominantly nuclear localization of DCHBV X likely leads to different functions compared with HBV X. Although it remains unclear how different localizations are determined, HBV(A) X and DCHBV(KT116) X may interact with host factors differently.

DDB2 induces the nuclear localization of HBV X without the binding of DDB1 (33). Because HBV X lacks a nuclear localization signal, its nuclear accumulation may rely on its interaction with other cellular proteins (34, 35). Therefore, it will be intriguing to test the molecular interaction of DCHBV X with DDB2 and other cellular proteins that also interact with HBV X.

In this study, we have identified two major isoforms of the feline *Smc6* gene. Isoform 1 has the KVRNT insertion, which is absent in other mammals, between amino acids 216 and 222 (**Figure 2**). Although our Smc6 degradation assay demonstrated that this insertion did not affect its sensitivity to mammalian hepatitis B virus X proteins (**Figures 6B and 6D**), the impact of this insertion on the antiviral activity should be examined. Interestingly, we have identified similar insertions encoding 1 to 3 amino acids in the human *Smc6* gene (rs1168792692, rs1669134078, and rs1669195959; https://www.ncbi.nlm.nih.gov/snp; accessed on 10/20/2024). The impact of these insertions, especially on the host’s susceptibility to HBV infection, can be investigated in future studies.

Mammalian hepatitis B virus X proteins have multiple functions in addition to Smc6 degradation. For example, HBV X modulates transcription, cell cycle control, cell growth, and apoptosis in hepatocytes (reviewed in 19, 36). Also, we have recently demonstrated that mammalian hepatitis B virus X proteins have conserved activities in preventing the TIR-domain-containing adaptor protein from inducing IFN-β (TRIF)-mediated IFN-β signaling pathway through TRIF degradation (37). We propose that mammalian hepatitis B virus X proteins exhibit different phenotypes depending on their functions and host species. It will be of interest to investigate which function is more and less conserved in mammalian hepatitis B virus X proteins. Furthermore, HBV X can be an attractive target for inhibitors (38).

There are several questions to be addressed in the future. First, we used an overexpression system to study Smc6 degradation by mammalian hepatitis B virus X proteins. It will be valuable to examine the roles of mammalian hepatitis B virus X proteins in Smc6 degradation with physiological protein levels. Second, the effect of the mammalian hepatitis B virus X proteins should be tested using an infectious and X-deficient strain of HBV to directly evaluate the roles of the mammalian hepatitis B virus X proteins in viral replication (25). Lastly, the domains or residues in HBV(A) X and DCHBV(KT116) X that determine different Smc6 degradation activities and DDB1 dependency can be investigated using chimeric proteins.

In summary, we have uncovered conserved yet distinct strategies employed by the mammalian virus of the genus *Orthohepadnavirus* in evading the host Smc5/6 defense system. Further research into the interaction between DCHBV(KT116) X, the Smc5/Smc6 complex, and related molecules can provide valuable insights into the molecular mechanisms underlying DCHBV pathogenesis. Understanding how DCHBV X targets and modulates the Smc5/6 complex may provide potential therapeutic insight into treating DCHBV-infected cats. Furthermore, it will contribute to the development of a novel HBV animal model using DCHBV.

## Materials and methods

### Cell culture

Lenti-X 293T cells (*Homo sapiens* or human; TaKaRa, Kusatsu, Japan, Cat# Z2180N), COS-7 (*Cercopithecus aethiops* or African green monkey [AGM]; Japanese Collection of Research Bioresources Cell Bank [JCRB], Ibaraki, Japan, Cat# JCRB9127), Fcwf-4 (*Felis catus* or cat; American Type Culture Collection, Manassas, VA, USA, Cat# CRL-2787), and CRFK (cat; JCRB, Cat# JCRB9035) cells were cultured in Dulbecco’s modified Eagle medium (DMEM; Nacalai Tesque, Kyoto, Japan, Cat# 08458-16) supplemented with 10% fetal bovine serum and 1 × penicillin-streptomycin (Nacalai Tesque, Cat# 09367-34).

### Plasmids

The cDNA sequences of the X protein derived from 9 viruses of *Orthohepadnavirus* were previously synthesized with codon optimization to humans (Twist Bioscience, San Francisco, CA, USA) (**Supplementary Table 2)** (37). cDNA fragments encoding X were cloned into a pCAGGS-mNeonGreen vector (39) predigested with AgeI-HF (New England Biolabs [NEB], Ipswich, MA, USA, Cat# R3552S) and NheI-HF (NEB, Cat# R3131M) using the In-Fusion Snap Assembly Master Mix (TaKaRa, Cat# Z8947N). The plasmids were amplified using NEB 5-alpha F′Iq Competent *Escherichia coli* (High Efficiency) (NEB, Cat# C2992H) and isolated using the PureYield Plasmid Miniprep System (Promega, Madison, WI, USA, Cat# A1222). The sequences of all the plasmids were verified using a SupreDye v3.1 Cycle Sequencing Kit (M&S TechnoSystems, Osaka, Japan, Cat# 063001) with a Spectrum Compact CE System (Promega).

### Smc5/6 complex degradation assay

Lenti-X 293T (1.25 × 10^5^ per well), COS-7 (4.17 × 10^4^ per well), CRFK (4.17 × 10^4^ per well), and Fcwf-4 cells (4.17 × 10^4^ per well) were plated in a 24-well plate (Thermo Fisher Scientific, Waltham, MA, USA, Cat# 142475). After overnight incubation, the cells were co-transfected with 500 ng of pCAGGS-mNeonGreen-X using the TransIT-X2 Dynamic Delivery System (TaKaRa, Cat# V6100) in the Opti-MEM I Reduced Serum Medium (Thermo Fisher Scientific, Cat# 31985062). After 48 h, the cells were stained with the NucBlue Live ReadyProbes Reagent (Hoechst 33342) (Thermo Fisher Scientific, Cat# R37605), observed under the EVOS M7000 Imaging System (Thermo Fisher Scientific), and collected for Western blotting analysis.

### Western blotting

Western blotting was used to examine Smc5/6 degradation by mammalian hepatitis B virus X proteins. First, pelleted cells were lysed in an M-PER Mammalian Protein Extraction Reagent (Thermo Fisher Scientific, Cat#78501) containing a Protease Inhibitor Cocktail Set I (×100) (Fujifilm, Osaka, Japan, Cat#165-26021). Protein concentrations were measured using the TaKaRa Bradford Protein Assay Kit (TaKaRa, Cat# T9310A). Then, 1 µg of the cellular extracts were mixed with 2cJ×cJBolt LDS sample buffer (Thermo Fisher Scientific, Cat# B0008) containing 2% β-mercaptoethanol (Bio-Rad, Hercules, CA, USA, Cat# 1610710) and incubated at 70°C for 10 min. The level of Smc6 was evaluated using a SimpleWestern Abby (ProteinSimple, San Jose, CA, USA) with an anti-SMC6L1 antibody (GeneTex, Irvine, CA, USA, Cat# GTX116832, ×50) and an Anti-Rabbit Detection Module (ProteinSimple, Cat# DM-001). The level of mNeonGreen-tagged X was measured with an anti-mNeonGreen antibody (Proteintech, Rosemont, IL, USA, Cat# 32F6-100, ×50) and an Anti-Mouse Detection Module (ProteinSimple, Cat# DM-002). The amount of input protein was measured using a Total Protein Detection Module (ProteinSimple, Cat# DM-TP01).

X protein localization was examined by first plating Lenti-X 293-T cells (1.25 × 10^5^ per well) in a 24-well plate. After overnight incubation, the cells were co-transfected with 500 ng of pCAGGS-mNeonGreen-X plasmids using the TransIT-X2 Dynamic Delivery System. After 48 h, the nuclear and cytosolic fractions were separated using a Minute Cytoplasmic and Nuclear Extraction Kit (Invent Biotechnologies, Inc., Cat# SC-003). The samples for Western blotting were prepared as described above. The localization of X was evaluated using an anti-mNeonGreen antibody. The efficiency of subcellular fractionation was evaluated using an anti-Lamin A/C antibody (Cell Signaling Technology [CST], Danvers, MA, USA, Cat# 4777S, ×250) and anti-α-Tubulin antibody (CST, Cat# 2125S, ×250).

### Smc6 degradation assay using the SpyTag/SpyCatcher system

First, total RNA was extracted from Lenti-X 293T, COS-7 CRFK, and Fcwf-4 cells using an RNeasy Mini Kit (QIAGEN, Chuo-ku, Japan, Cat# 74104) and QIAshredder (QIAGEN, Cat# 79656). Then, *Smc6* cDNA was amplified by RT-PCR using the PrimeScript II High Fidelity One Step RT-PCR Kit (TaKaRa, Cat#R026B) using specific primers (**Supplementary Table 3**). The PCR consisted of 1 cycle of 45°C for 10 min; 1 cycle of 94°C for 2 min; 40 cycles of 98°C for 10 s, 60°C for 15 s, and 68°C for 20 s; 1 cycle of 68°C for 7 min. The amplified fragments were ligated into pCAGGS-SpyTag-sfCherry3C(11)-HA predigested with AgeI-HF and NheI-HF using an In-Fusion Snap Assembly Master Mix. The plasmids were verified by sequencing.

The system was validated by plating Lenti-X 293T cells at 2.5cJ×cJ10^5^ cells per well in a 12-well plate. Then the cells were transfected with pCAGGS-SpyCatcher-sfCherry3C(1–10) alone, pCAGGS-SpyTag-sfCherry3C(11)-mNeonGreen alone, or a combination of pCAGGS-SpyCatcher-sfCherry3C(1–10), pCAGGS-SpyTag-sfCherry3C(11)-mNeonGreen, and pCAGGS-SpyTag-sfCherry3C(11)-HA-Smc6 (Human). The signals of mNeonGreen and sfCherry3C were measured 2 days after transfection using an Attune NxT Flow Cytometer (Thermo Fisher Scientific). The mNeonGreen- and sfCherry3C-double-positive population was gated and analyzed using FlowJo v10.8.1 (Becton, Dickinson and Company).

To evaluate the role of DDB1 in Smc6 degradation, we depleted human DDB1 by transfecting Lenti-X 293T cells (1.9cJ×cJ10^6^ cells per well in a 6-well plate) with DsiRNA targeting human *Ddb1* (Integrated DNA Technologies [IDT], Coralville, IA, USA, Reference# hs. Ri. DDB1.13.1, and hs. Ri. DDB1.13.3) or Negative Control DsiRNA (IDT, Cat# 109617253) with the TransIT-X2 Dynamic Delivery System. We depleted feline Ddb1 by transfecting Fcwf-4 cells (6.3cJ×cJ10^5^ cells per well in a 6-well plate) with DsiRNA targeting feline *Ddb1* (IDT, Reference# CD.Ri.477260.13.1 and CD.Ri.477260.13.3) or Negative Control DsiRNA with the TransIT-X2 Dynamic Delivery System. After 2 days, the cells were re-plated in a new 96-well plate at 3cJ×cJ10^4^ Lenti-X 293T cells per well or 1cJ×cJ10^4^ Fcwf-4 cells per well. The cells were cultured overnight and co-transfected with pCAGGS-SpyCatcher-sfCherry3C(1–10) at 15 ng/well, pCAGGS-SpyTag-sfCherry3C(11)-HA-Smc6 at 15 ng/well, and pCAGGS-mNeonGreen-X at 70 ng/well. Smc6 degradation was measured with sfCherry3C-positivity 2 days after transfection as described above.

### Alignment of the mammalian hepatitis B virus X proteins and phylogenetic analysis

The complete amino acid sequences of the X protein from 32 virus strains of the genus *Orthohepadnavirus* were aligned using the MUSCLE algorithm in MEGA X (MEGA Software). The alignment parameters were gap open at −2.90, gap extend at 0.00, and hydrophobicity multiplier at 1.20. Alignment visualization was done using CLC Genomics Workbench viewing mode Version 22.0.1.

A phylogenetic tree was built using the alignment of X amino acid sequences retrieved from public databases, and evolutionary analysis was conducted using MEGA X (40, 41). The evolutionary history was inferred using the maximum likelihood method and the JTT matrix–based mode (42). The initial trees for the heuristic search were obtained by applying the neighbor-joining method to a matrix of pairwise distances estimated using the JJT model. A discrete gamma distribution was used to model the evolutionary rate differences among sites. The tree is drawn to scale, with branch lengths measured in the number of substitutions per site.

### Alignment of the mammalian Smc6 proteins and phylogenetic analysis

The regions around residues 36 to 48 and 216 to 222 of the Smc6 protein from 23 animal species and 3 human cells were aligned using the MUSCLE algorithm in MEGA X (MEGA Software). The alignment parameters were gap open at −2.90, gap extend at 0.00, and hydrophobicity multiplier at 1.20.

A phylogenetic tree was constructed using the alignment of Smc6 amino acid sequences retrieved from public databases. The evolutionary analysis was conducted in MEGA X. The evolutionary history was inferred using the maximum likelihood method and the JTT matrix–based mode. The initial trees for the heuristic search were obtained by applying the neighbor-joining method to a matrix of pairwise distances estimated using the JJT model. A discrete gamma distribution was used to model the evolutionary rate differences among sites. The tree is drawn to scale, with branch lengths measured in the number of substitutions per site.

### Calculation of the identity of Smc6 among animal species

The identity of Smc6 among the animal species was calculated using MEGA X with a pairwise distance matrix. Analyses were conducted using the JTT matrix-based model (43). The analysis involved 26 amino acid sequences. All ambiguous positions were removed for each sequence pair (pairwise deletion option). There were a total of 1125 positions in the final dataset. Evolutionary analyses were conducted in MEGA11 (44, 45).

### Statistical analysis

The level of Smc6 degradation was evaluated by one-way ANOVA, followed by the Tukey test. A *p*-value of 0.05 or less was considered statistically significant. The analysis was performed using Prism 10 v10.2.3 (GraphPad Software).

## Acknowledgments

The authors thank Ms. Tomoko Nishiuchi, Ms. Yuki Shibatani, Ms. Miki Kawano, Ms. Natsumi Matsubara, and the staff of CADIC, University of Miyazaki, for their assistance for their assistance. Supplementary Figure 1 was created with BioRender (https://biorender.com/).

## Author information Contributions

M.S. and A.S. designed the experiments. M.S., YV.F, and A.S. performed the experiments. M.S., YV.F, and A.S. analyzed the results. M.S. and A.S. wrote the manuscript. All authors have read and approved the manuscript.

## Data availability

Source data are available on request.

## Ethics declarations

N/A

---

**Supplemental Figure 1.** Schematic diagram of the SpyTag/SpyCatcher-assisted SMC6 degradation assay. Flow cytometry analysis of the fluorescence in cells producing SpyCatcher-sfCherry3C(1–10) and SpyTag-sfCherry3C(11)-HA-Smc6. In the absence of mammalian hepatitis B virus X proteins, a specific fluorescence signal was detected when SpyCatcher-sfCherry3C(1–10) was reconstituted with the SpyTag-sfCherry3C(11)-HA-Smc6 to generate a functional sfCherry3C protein (upper panel). In the presence of mammalian hepatitis B virus X proteins, the signal was decreased due to the degradation of SpyTag-sfCherry3C(11)-HA-Smc6 by X proteins (lower panel).

