## Supplementary Table for "Conserved yet Divergent Smc5/6 Complex Degradation by Mammalian Hepatitis B Virus X Proteins"

### Supplementary Table 1. Genetic distance of mammalian Smc6 based on amino acid sequence

The number of amino acid substitutions per site from between sequences are shown.

| No | | 1 | 2 | 3 | 4 | 5 | 6 | 7 | 8 | 9 | 10 | 11 | 12 | 13 | 14 | 15 | 16 | 17 | 18 | 19 | 20 | 21 | 22 | 23 | 24 | 25 | 26 |
|---|---|---|---|---|---|---|---|---|---|---|---|---|---|---|---|---|---|---|---|---|---|---|---|---|---|---|---|
| 1 | NP_001135758.1_Homo_sapiens | | | | | | | | | | | | | | | | | | | | | | | | | | |
| 2 | AWK27963.1_Homo_sapiens_293T | 0.00 | | | | | | | | | | | | | | | | | | | | | | | | | |
| 3 | AWK27960.1_Homo_sapiens_HepG2 | 0.00 | 0.00 | | | | | | | | | | | | | | | | | | | | | | | | |
| 4 | XP_054404336.1_Pongo_abelii | 0.01 | 0.01 | 0.01 | | | | | | | | | | | | | | | | | | | | | | | |
| 5 | XP_055234727.1_Gorilla_gorilla_gorilla | 0.00 | 0.01 | 0.00 | 0.01 | | | | | | | | | | | | | | | | | | | | | | |
| 6 | AWK27961.1_Chlorocebus_tantalus_Cos-7 | 0.02 | 0.02 | 0.02 | 0.02 | 0.02 | | | | | | | | | | | | | | | | | | | | | |
| 7 | AWK27957.1_Chlorocebus_pygerythrus_Vero | 0.02 | 0.02 | 0.02 | 0.02 | 0.02 | 0.00 | | | | | | | | | | | | | | | | | | | | |
| 8 | XP_007969584.1_Chlorocebus_sabaeus | 0.02 | 0.02 | 0.02 | 0.02 | 0.02 | 0.00 | 0.00 | | | | | | | | | | | | | | | | | | | |
| 9 | XP_005576553.2_Macaca_fascicularis | 0.02 | 0.02 | 0.02 | 0.02 | 0.02 | 0.00 | 0.00 | 0.00 | | | | | | | | | | | | | | | | | | |
| 10 | XP_014967240.1_Macaca_mulatta | 0.02 | 0.02 | 0.02 | 0.02 | 0.02 | 0.00 | 0.00 | 0.00 | 0.00 | | | | | | | | | | | | | | | | | |
| 11 | AWK27958.1_Aotus_trivirgatus_OMK | 0.04 | 0.04 | 0.04 | 0.04 | 0.04 | 0.04 | 0.04 | 0.04 | 0.04 | 0.04 | | | | | | | | | | | | | | | | |
| 12 | AWK27962.1_Saguinus_oedipus_B95a | 0.04 | 0.04 | 0.04 | 0.04 | 0.04 | 0.04 | 0.04 | 0.04 | 0.04 | 0.04 | 0.02 | | | | | | | | | | | | | | | |
| 13 | AWK27959.1_Mustela_putorius_MPF | 0.06 | 0.06 | 0.06 | 0.06 | 0.06 | 0.06 | 0.06 | 0.06 | 0.06 | 0.06 | 0.08 | 0.08 | | | | | | | | | | | | | | |
| 14 | XP_023107850.1_Felis_catus_Isoform_X1 | 0.08 | 0.07 | 0.08 | 0.08 | 0.08 | 0.08 | 0.08 | 0.08 | 0.07 | 0.07 | 0.08 | 0.08 | 0.04 | | | | | | | | | | | | | |
| 15 | XP_003984536.2_Felis_catus_isoform_X2 | 0.08 | 0.07 | 0.08 | 0.08 | 0.08 | 0.08 | 0.08 | 0.08 | 0.07 | 0.07 | 0.08 | 0.08 | 0.04 | 0.00 | | | | | | | | | | | | |
| 16 | XP_038416754.1_Canis_lupus_familiaris | 0.07 | 0.07 | 0.07 | 0.07 | 0.07 | 0.07 | 0.07 | 0.07 | 0.07 | 0.07 | 0.07 | 0.08 | 0.03 | 0.01 | 0.01 | | | | | | | | | | | |
| 17 | XP_027629216.1_Tupaia_chinensis | 0.07 | 0.07 | 0.07 | 0.07 | 0.07 | 0.07 | 0.07 | 0.07 | 0.07 | 0.07 | 0.07 | 0.07 | 0.08 | 0.07 | 0.07 | 0.06 | | | | | | | | | | |
| 18 | XP_038968116.1_Rattus_norvegicus | 0.09 | 0.09 | 0.09 | 0.09 | 0.10 | 0.10 | 0.10 | 0.10 | 0.09 | 0.10 | 0.10 | 0.10 | 0.10 | 0.09 | 0.09 | 0.08 | 0.09 | | | | | | | | | |
| 19 | NP_079971.2_Mus_musculus | 0.10 | 0.10 | 0.10 | 0.10 | 0.10 | 0.10 | 0.10 | 0.10 | 0.10 | 0.10 | 0.10 | 0.10 | 0.11 | 0.09 | 0.09 | 0.09 | 0.09 | 0.02 | | | | | | | | |
| 20 | XP_002691521.2_Bos_taurus | 0.08 | 0.08 | 0.08 | 0.08 | 0.09 | 0.09 | 0.09 | 0.09 | 0.08 | 0.08 | 0.09 | 0.09 | 0.08 | 0.07 | 0.07 | 0.06 | 0.07 | 0.10 | 0.10 | | | | | | | |
| 21 | XP_003125390.1_Sus_scrofa | 0.08 | 0.09 | 0.08 | 0.08 | 0.09 | 0.08 | 0.08 | 0.08 | 0.08 | 0.08 | 0.08 | 0.09 | 0.08 | 0.07 | 0.07 | 0.06 | 0.07 | 0.10 | 0.11 | 0.06 | | | | | | |
| 22 | XP_032980470.1_Rhinolophus_ferrumequinum | 0.09 | 0.09 | 0.09 | 0.09 | 0.09 | 0.09 | 0.09 | 0.09 | 0.09 | 0.09 | 0.09 | 0.09 | 0.09 | 0.08 | 0.08 | 0.07 | 0.08 | 0.10 | 0.11 | 0.08 | 0.08 | | | | | |
| 23 | XP_016009374.1_Rousettus_aegyptiacus | 0.09 | 0.09 | 0.09 | 0.09 | 0.10 | 0.09 | 0.09 | 0.09 | 0.08 | 0.09 | 0.09 | 0.09 | 0.09 | 0.08 | 0.08 | 0.08 | 0.08 | 0.11 | 0.11 | 0.09 | 0.08 | 0.06 | | | | |
| 24 | XP_006910309.1_Pteropus_alecto | 0.09 | 0.09 | 0.09 | 0.09 | 0.09 | 0.09 | 0.09 | 0.09 | 0.08 | 0.08 | 0.09 | 0.09 | 0.08 | 0.07 | 0.07 | 0.07 | 0.07 | 0.10 | 0.10 | 0.08 | 0.08 | 0.06 | 0.03 | | | |
| 25 | XP_023384272.1_Pteropus_vampyrus | 0.10 | 0.10 | 0.10 | 0.10 | 0.10 | 0.10 | 0.10 | 0.10 | 0.09 | 0.10 | 0.10 | 0.10 | 0.09 | 0.08 | 0.08 | 0.08 | 0.09 | 0.11 | 0.12 | 0.09 | 0.09 | 0.08 | 0.04 | 0.01 | | |
| 26 | XP_028932004.1_Ornithorhynchus_anatinus | 0.39 | 0.39 | 0.39 | 0.39 | 0.40 | 0.39 | 0.39 | 0.39 | 0.39 | 0.39 | 0.40 | 0.40 | 0.39 | 0.39 | 0.39 | 0.39 | 0.39 | 0.41 | 0.41 | 0.40 | 0.40 | 0.39 | 0.40 | 0.41 | 0.43 | |

**Supplementary Table 2. Synthesized DNAs for generating plasmids encoding X protein**

| X protein | Codon-optimized DNA sequence |
|---|---|
| HBV(genotype A)<br>Accession# LC488828.<br>Host: human | ATGTACCCTTACGATGTACCTGACTACGCGACCGGTGCCGCT<br>CGACTTTATTGCCAACTTGATCCATCCCGCGACGTGTTGTGT<br>CTCAGACCCGTTGGCGCCGAATCTCGAGGTCGCCCACTTTCA<br>GGACCCCTTGGTACATTGAGTTCTCCATCTCCGAGCGCTGTA<br>CCGGCGGACCACGGGGCTCATTTGAGTTTGAGGGGTCTCCC<br>AGTGTGTGCTTTCAGCAGTGCCGGCCCGTGCGCGCTTCGATT<br>CACTTCAGCAAGATGTATGGCTACGACTGTCAATGCTCATCA<br>AATCCTCCCAAAGGTCCTCCACAAAAGAACGCTTGGGCTTCC<br>GGCCATGTCTACTACTGATCTTGAGGCGTACTTCAAGGACTG<br>CGTCTTTAAGGATTGGGAGGAATTGGGGGAGGAAATTCGCC<br>TTAAAGTATTCGTGCTGGGGGGTTGCAGACACAAACTTGTAT<br>GCGCTCCTGCCCCGTGCAACTTCTTCACTTCCGCATAA |
| HBV(genotype D)<br>Accession# YP_009173867.1<br>Host: human | ATGTACCCTTACGATGTACCTGACTACGCGACCGGTGCCGCA<br>CGCCTCTGTTGCCAACTTGATCCAGCACGGGATGTTCTGTGC<br>CTTCGGCCAGTCGGCGCTGAAAGTTGTGGACGGCCGTTCTCC<br>GGCTCCCTCGGAACGCTCTCCTCCCCCTCCCCCTCAGCCGTC<br>CCTACAGATCATGGAGCTCACCTTTCTCTTCGCGGGCTTCCG<br>GTATGCGCGTTTTCTTCTGCTGGACCTTGCGCGTTGCGCTTTA<br>CATCTGCTAGGAGAATGGAAACAACGGTTAACGCGCACCAG<br>ATACTCCCGAAGGTTTTGCACAAAAGGACCCTGGGTCTTAGT<br>GCCATGAGCACTACTGATCTCGAGGCATACTTTAAGGACTGC<br>TTGTTCAAGGATTGGGAGGAACTTGGGGAAGAGATCAGATT<br>GAAGGTGTTCGTCTTGGGGGGATGTCGGCATAAGCTCGTGT<br>GTGCACCTGCACCCTGCAATTTCTTCACGTCTGCTTAG |
| HBV(genotype G)<br>Accession# BAD91282.1<br>Host: human | ATGTACCCTTACGATGTACCTGACTACGCGACCGGTGCAGCC<br>CGGCTCTGTTGTCAGTTGGATCCCAGCCGAGACGTTCTCTGC<br>TTGCGGCCAGTTAGCGCAGAGTCATCTGGACGCCCGCTGCCC<br>GGACCTTTTGGTGCACTCTCTCCTCCAAGTCCCTCTGCAGTC<br>CCCGCAGATCATGGCGCTCACCTGTCATTGCGCGGTTTGCCA<br>GTATGCGCCTTCTCCTCAGCAGGTCCGTGCGCGCTTCGATTC<br>ACGTCAGCACGGTACATGGAAACGGCAATGAATACGAGCCA<br>CCATCTCCCCCGACAACTTTACAAACGCACCCTCGGTCTTTT<br>CGTTATGAGTACCACTGGTGTCGAGAAGTACTTTAAAGACTG<br>TGTCTTCGCTGAATGGGAGGAGTTGGGCAACGAGAGCCGAC<br>TGATGACTTTCGTGCTGGGCGGATGCAGACATAAGCTCGTGT<br>GTGCTCCGGCCCCCTGTAATTTCTTTACTTCAGCTTAA |
| HBV(genotype H)<br>Accession# BAJ15483.1<br>Host: human | ATGTACCCTTACGATGTACCTGACTACGCGACCGGTGCCGCT<br>AGGCTGTGCTGCCAACTTGATCCCGCCCGCGACGTGCTGTGT<br>TTGCGGCCAGTAGGTGCTGAAAGCTGCGGTCGGCCACTTTCC<br>TGGTCCCTCGGGGCTTTGCCCCCGTCATCACCTCCAACAGTT<br>CCTGCTGACGACGGATCTCACTTGAGTCTTCGGGGGTTGCCC<br>GCGTGCGCGTTCAGTTCCGCTGGTCCTTGTGCGTTGCGATTT<br>ACGAGTGCTAGGAGAATGGAAACGACTGTCAACGCTCCCTG<br>GAACTTGCCCACTACGCTGCACAAACGAACATTGGGTCTGTC<br>TCCCCGCTCCACGACTTGGATAGAAGAGTACATTAAAGACT<br>GTGTTTTCAAAGACTGGGAGGAAAGTGGGGAGGAACTTCGC<br>CTGAAGGTGTTTGTGCTTGGCGGTTGTAGACACAAACTCGTG<br>TGTTCCCCGGCCCCTTGTAACTTCTTCACTAGCGCATAG |

| | |
|---|---|
| DCHBV(KT116)<br>Accession# LC668427.1<br>Host: Domestic cat | ATGTACCCTTACGATGTACCTGACTACGCGACCGGTGCAGCA<br>CGGCTGCGCTGCGAACTCGATCCTTCTGGTCGGGTCCTGCGG<br>TTGAGACCATTCATTAGTGAATCCAGCGGACGCGCGGTTAG<br>CCGAACTGCACGCTTGCCAGACCTGAGCCCCTCAGTTGCGGT<br>TTCAGCGACACTGCGGGCCAGGGAATCTCTGAGGGGTATAC<br>CTGCCTGTCTCACGTCACCAGAGGGCCCTTGTGTTTTGAGAT<br>TTACCTGCGCTGATAGTAGACGGTGCATGGAAGCAGCAATG<br>ATTGGCTTGGTCCCAGCACTGCTTGCTCGCCAACTTGGCTTC<br>GGGACTTGGCAGCCGGATGTATGGACGCTTCGGCTTCGCGA<br>TCTTTTGTTGGTCGAGTGGGAGGAAGAAGGACTGACGCCGC<br>GGTTGTGTACTTATCTTGTAACGGGGTGCGCTCATAAAACGC<br>TTCACACTCGATAG |
| Domestic donkey HBV<br>Accession# QMV34684.1<br>Host: Domestic donkey | ATGTACCCTTACGATGTACCTGACTACGCGACCGGTGCGGCT<br>CGCCTGAGGTGTCAACTGGACCCAACTGGCCGGGTACTCCA<br>CCTCCGACCCTTTACTTCCGAATCTTGTAGAAGAACTTTGGC<br>AGGTACTGCCGGGGCACCAGATCTCCCAGCAGCGGACCTCC<br>TTCAAGCGGATCACCGGACTCATCTTAGGGTTCGACGCTTGC<br>CTGCTTGCTGTTTCTCTTCTCGCGGTCCGTGTGTGCTTAGGTT<br>CACATGCGCGGACCTTAGCCGACGAATGGAAGCCCCGATGA<br>ACCTCGTTCAATATCTGGGGAAAAGGGCGCGGGGTCTTCAG<br>CATCCGCCCGGTGATTCCTATTGCCAACATGAACTTTGGACA<br>CAATGGGAGGAGAATGGTTGGTCAGACAGAATCTATACTTA<br>CGTGTTGGGAGGATGCAGACACAAATGGCTTTACCCACTTTA<br>G |
| Asian grey shrew HBV<br>Accession# YP_010796421.1<br>Host: Asian grey shrew | ATGTACCCTTACGATGTACCTGACTACGCGACCGGTGCTGCA<br>AGAATGCTCTTCGATCTCGACCCTGCTACAGGAGCTGTACGC<br>CTTCGCCCATTTCTCACTGAACCCCGCGGACGAGGGGAACA<br>GACACCGCGCCCGACTTCCTCTCCGACAACGTCAGCCCTGTC<br>TTCTTTCCTTGGAAGTCGGTCTTCCTGGCGGCGCTTGCCAAG<br>CTGCGCCGACTCTCCATTCGGCCCATGTACTTTGCGGTTCAC<br>GTTCGCAGAGCTGGGAAACTTGCAGACACCAATGAACTCAG<br>TGACCTTCATCAGTTGTCGGTCAAGGGGAGCCCATCTGAAGT<br>GCCGGAGGCAACAGAAGAATTGGACCTGGTATTTCTGGACA<br>CATCATAATGCGAACAACACGCACCATTTGTGGCTTATGTGC<br>TACGGAGGTTGTAGGCATAAATAG |
| Capuchin monkey HBV<br>Accession# YP_009666527.1<br>Host: Capuchin monkey | ATGTACCCTTACGATGTACCTGACTACGCGACCGGTGCAGCC<br>AGACTTTGTTGCCAACTGGACCCTGCCAGGGATGTTCTTTGT<br>CTCCGACCTGTAACTGCCGAGCCATGTGGACGACCCTTCAGC<br>GGTTCTGCTCGGACATCCGCTCCGGCAGCTGCGGCAGCCCTG<br>CCCTCTATTGATGGAGCATATCTGTCCCTTCGAGGGCTTCCT<br>AGTTGCGCTTTCTCATCCTCAGGGCCCTGCGCCTTGAGGTTT<br>ACAAGTGCGCGACGAATGGCTACACCGATGAATAGTAGAGA<br>TCTGGTCCAACAACTCTATAATCGGACGTTGGGTCTTGCTCC<br>TCTCTCCACTGGGCAGTGGGAACGGCACTTTAAAGATCTTTT<br>GTTCGAGGAATGGGAGGAACTCGGTGTTGAGTTCAGGTTGA<br>AAGTATTCGTGCTGGGGGGTTGTCGCCATAAGCTCGTTTGCA<br>GTGTGCAACCTTGCATATTCTTCACTAGTGCCTAA |
| Orangutan HBV<br>Accession# AAF33122.1<br>Host: Orangutan | ATGTACCCTTACGATGTACCTGACTACGCGACCGGTGCTGCC<br>CGCCTTTGTTGTCAGTTGGACCCGGCTCGAGATGTCCTTTGC<br>CTTCGGCCAGTGGGAGCTGAGAGTAGAGGAAGGCCGTTCCC<br>AGGCAGTATTGGTGCTCTGCCCCCACCATCTCTGAGCGCGGT<br>ACCGGCCGACCACGGAGCCCACCTTAGCCTTCGGGGTTTGC<br>CGGTATGCGCTTTTTCTTCAGCGGGGCCTTGTGCGTTGAGGT |

| | |
|---|---|
| | TCACGAGCGCCAGGTGCATGGAGACCACCGTAAATGCGCCT<br>AGAAATCTCCCTAAGGTCCTGCATAAGAGAACATTGGGCCT<br>TTCCACTATGTCTACTACGCGAATCGAAACGTACTTTAAGGA<br>TTGTGTGTTTAAGGATTGGGAAGAACTTGGAGAGGAGATCC<br>GGTTGAAGGTTTTTGTCTTGGGTGGATGTAGGCATAAATTGG<br>TGTGTTCTCCCGCGCCTTGCAACTTTTTTACAAGTGCATGA |

**Supplementary Table 3. Primers used for generating a plasmid encoding Smc6 protein**

| Host | Direction | Sequence (5′-3′) |
|---|---|---|
| Human | Forward | TGACTACGCGACCGGTGCCAAAAGAAAGGAAGAAAATTTT |
| | Reverse | AAAAAGATCTGCTAGCTCACCTTTGGTCATCATCTTCTTCT |
| Feline | Forward | GATGTACCTGACTACGCGACCGGTGCCAAAAGAAAGGAAGAAAA |
| | Reverse | AAAAAGATCTGCTAGCTCAGCTCCGGTCTTCTTCCTCC |
| Feline (+KVRNT) | Forward | AAAGTAAGGAACACCAAATTCTTTATGAAAGCAAC |
| | Reverse | GGTGTTCCTTACTTTGTATTTGTCTCCCTCATTTT |
