## Supplementary figures and images for "Conserved yet Divergent Smc5/6 Complex Degradation by Mammalian Hepatitis B Virus X Proteins"

### Supplementary Figure

#### Supplemental Figure 1

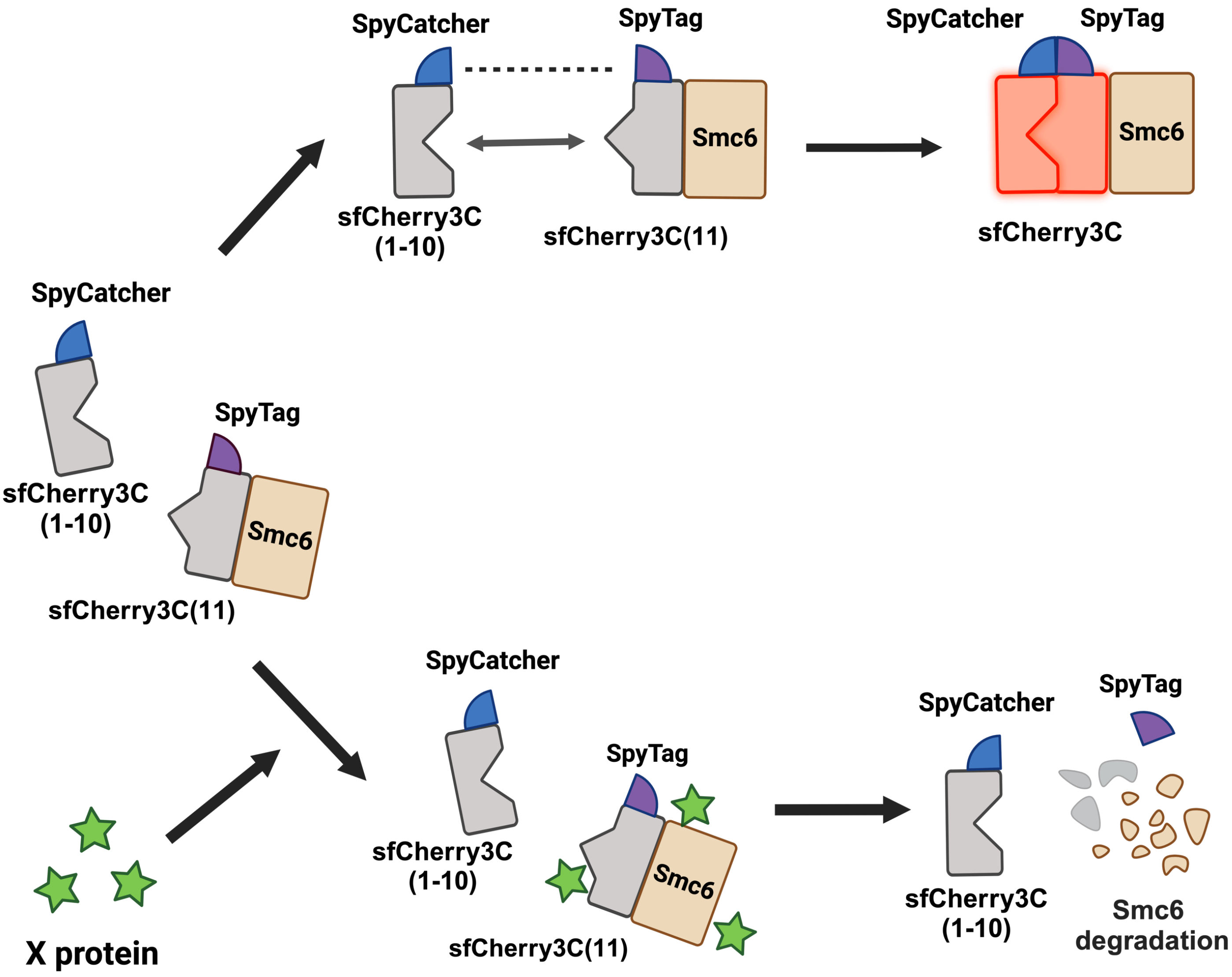
